# Sex differences in the cardiopulmonary and neuromuscular response to high-intensity interval exercise

**DOI:** 10.1101/2024.11.08.622119

**Authors:** Hannah Wilson, Lina Bernert, Padraig Spillane, Emma Squires, Lorna Crawford, Jessica Piasecki, Ross Julian, Eurico N. Wilhelm, Kirsty M Hicks, Paul Ansdell

**Author notes:** Corresponding author: Paul Ansdell Ph.D, FHEA, Department of Sport, Exercise and Rehabilitation Faculty of Health and Life Sciences, Northumbria University Newcastle upon Tyne Tyne and Wear, NE1 8ST, United Kingdom. Equal contribution to the manuscript (Hannah Wilson, Lina Bernert).

## Abstract

Sex differences exist in the integrative response to exercise, however, these are typically researched during constant-load exercise. Interval exercise involves high-intensity efforts interspersed with recovery periods to repeatedly stress physiological systems, and it is currently unknown whether the response to this form of exercise differs between sexes.

Ten males and ten females (age: 25±3 years) completed two experimental visits. First, an incremental treadmill exercise test was performed to obtain submaximal (lactate threshold) and maximal (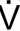
O_2peak_) data. Thereafter, visit two involved 4 × 3-min running intervals at 90% of the final incremental test velocity (v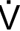
O_2peak_), with 90 secs rest between intervals. Before exercise and after each interval, maximal voluntary contraction (MVC), quadriceps potentiated twitch (Q_tw.pot_), and voluntary activation (VA) were recorded. The rates of oxygen uptake (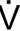
O_2_), carbon dioxide production (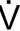
CO_2_) and ventilation (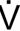
_E_) were continuously recorded throughout.

There was no sex difference in relative 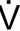
O_2peak_ (males: 47.2±6.0 vs. females: 44.4±5.8 ml.kg^-1.^min^-1^, p=0.292). When expressed relative to peak values, there were no sex differences in the 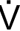
O_2_ or 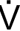
CO_2_ response to the interval task (p≥0.781). Females had greater 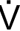
_E_, 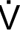
_E_/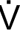
O_2_, and 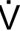
_E_/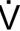
CO_2_ values during the first two intervals (p≤0.046). There were no sex differences in the reductions in MVC, Q_tw.pot_, and VA during the interval task (p≥0.150), however females had lesser reductions in Q_tw.pot_ values post-exercise (−24±9 vs. −15±8%, p=0.044).

Sex differences exist in the physiological response to interval exercise. Compared to males, females experienced greater hyperpnoea during the initial stages, and had lesser decreases in contractile function post-exercise.

## Introduction

Females have historically been under-represented in sport science studies for a multitude of reasons (Cowley *et al*., 2021; James *et al*., 2023). As a result, assumptions about exercise training have been generalised from male-dominated research and applied to females. It is becoming increasingly evident that the acute physiological responses to various modalities of exercise differ between the sexes (Ansdell *et al*., 2020b), meaning this approach might not be optimal for the prescription of exercise to females. Research is needed to identify whether males and females respond to commonly prescribed forms of exercise similarly, in order to optimise training and performance for both sexes.

Morphological and anatomical sex differences in key physiological systems are thought to lead to the differences in the integrative response to exercise between sexes. For instance, males typically have a greater quantity of muscle mass and can generate larger maximal force, but experience greater proportional occlusion of limb blood flow during muscle contraction (Hammer *et al*., 2023). Females have consistently demonstrated to have a higher proportional area of type I muscle fibres (Staron *et al*., 2000), as well as greater capillary density (Roepstorff *et al*., 2006), and vasodilatory response of the femoral artery (Parker *et al*., 2007). These physiological sex differences have previously been suggested to aid in oxygen delivery and help delay the onset of fatigue in females (Hunter, 2014; Ansdell *et al*., 2020b). Despite the potentially superior aerobic muscular phenotype, it has been established that females have smaller lung volumes, airway size, and alveolar surface for gas exchange (Mead, 1980; Crapo *et al*., 1982; Martin *et al*., 1987; Guenette *et al*., 2009). These morphological differences lead to a greater work and oxygen cost of breathing at elevated ventilatory rates (Guenette *et al*., 2009; Sheel *et al*., 2016), resulting in an increased fraction of whole-body oxygen uptake (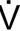
O_2_) originating from the respiratory musculature in females compared to males (Dominelli *et al*., 2015). This, combined with lower haemoglobin concentrations (Murphy, 2014), means that females have a poorer O_2_ carrying capacity during exercise (Harms *et al*., 1998; Diaz-Canestro *et al*., 2022). The balance between the importance of O_2_ delivery and utilisation depends on the physiological determinants of the task being performed, which is one reason why sex differences in the integrative response to exercise are not uniform and require further investigation (Hunter, 2016; Ansdell *et al*., 2020b).

Previous studies considering single-limb contractions found that males experienced greater rates of fatigue than females (Hunter *et al*., 2004; Ansdell *et al*., 2017; Senefeld *et al*., 2018) as well as males demonstrating a slower recovery time than females (Senefeld *et al*., 2018; Ansdell *et al*., 2019a). Indeed, more recent evidence suggests that these sex differences in the response to single-limb exercise appear to be related to the proportion of myosin heavy chain I isoform and mitochondrial protein abundance (McDougall *et al*., 2023). Despite this evidence in single-limb models, these findings do not necessarily translate into whole-body locomotion (Sidhu *et al*., 2013). In particular, the literature comparing the physiological response to locomotor exercise between sexes is less comprehensive. Several studies have demonstrated that female knee-extensors experience less fatigue following cycling to task failure at equivalent relative exercise intensities (Ansdell *et al*., 2020a; Azevedo *et al*., 2021; Azevedo *et al*., 2022), or following fixed duration running (Glace *et al*., 1998) and cycling (Glace *et al*., 2013) tasks. Although these studies point towards a consistent mechanism for the sex difference in fatigability, the tasks employed are limited in generalisability to athletic training outside of the controlled lab environment. Constant-load tasks, particularly to task failure, are rarely employed by those prescribing exercise for athletic enhancement. Recently, we demonstrated that the sex difference in fatigability was evident following a self-paced 5 km running time trial (Solleiro Pons *et al*., 2023), whilst others have demonstrated similar sex differences following longer distance running tasks longer than 40 km (Temesi *et al*., 2015; Besson *et al*., 2021). Although Boccia *et al*. (2018) did not observe a sex difference in fatigability following a half marathon, suggesting that the difference is intensity and/or duration dependent.

Interval exercise is a training method often prescribed to enhance aerobic and anaerobic capacity (Laursen & Jenkins, 2002). Research that has previously investigated sex differences in the response to interval exercise typically utilises repeated sprints (5-6 secs bouts) interspersed with prolonged (25-30 secs) rest (Brooks *et al*., 1990; Billaut & Smith, 2009). This modality of training has implications for intermittent and team sport athletes, and researchers have previously suggested that the lesser fatigue experienced by females is related to lower absolute mechanical work (Billaut & Bishop, 2012). Longer duration intervals are typically prescribed at submaximal, yet supra-threshold intensities (e.g., 2-6 min at 3 km-10 km race pace, Parmar *et al.,* 2021). In terms of exercise intensity domains, the intention of such intervals is to intersperse severe intensity bouts with periods of moderate intensity recovery or rest, and repeatedly elicit a state of metabolic stress (Jones & Vanhatalo, 2017). Given that evidence from constant-load exercise indicates that females are more resistant to fatigue during metabolically challenging tasks (Ansdell *et al*., 2020a; Azevedo *et al*., 2021; Azevedo *et al*., 2022), it is important to understand how interspersing high-intensity bouts with recovery periods mediates the sex difference in fatigability and the integrative physiological response to exercise. The little evidence that does exist from cycling exercise suggests that there might be perceptual, but not physiological sex differences in the response to high-intensity interval exercise (Tripp *et al*., 2024). As highlighted, both exercise modality and intensity mediate the influence of sex on the responses to exercise, therefore this study aimed to compare the physiological responses to, and recovery from, a bout of high-intensity interval running between sexes. We hypothesised that both sexes would experience a similar metabolic response to exercise, but females would be less fatigable.

## Methods

### Ethical Approval

This study received institutional ethical approval from the Northumbria University Health and Life Sciences Research Ethics Committee (submission reference: 2022-0094-368) and was conducted according to all aspects of the Declaration of Helsinki, apart from pre-registration in a public database. Participants volunteered for the study and provided written informed consent.

### Sample Size Calculation

An *a priori* sample size calculation was performed using GPower (v3.0.0) using the effect size from Ansdell *et al*. (2020a) for the sex difference in contractile dysfunction following high-intensity cycling (ηp² = 0.344), and reliability data for the same variable from Ansdell *et al*. (2019b). With the parameters of α = 0.001 and 1-β = 0.99, the minimum sample size required was 12 participants (6 males, 6 females). Therefore, to maximise statistical power, 10 participants of each sex were recruited.

### Participant Characteristics

Ten healthy males (mean ± SD age: 27 ± 4 years, stature: 180 ± 6 cm, body mass: 83.5 ± 12.1 kg) and ten healthy females (age: 23 ± 2, stature: 163 ± 6, body mass: 62.9 ± 9.1 kg) whose gender matched their sex assigned at birth, volunteered for the study. All participants were free from musculoskeletal conditions, as well as neurological, respiratory, and cardiovascular disease. Hormonal status was not an exclusion criterion or controlled for in this study, based on evidence that the menstrual cycle or hormonal contraceptive usage do not influence the cardiopulmonary or neuromuscular responses to whole-body exercise (Georgescu *et al*., 2020; Mattu *et al*., 2020). Five females were naturally cycling, with their second visits occurring on self-reported days 5, 8, 16, 19, and 23 of their menstrual cycle. Three females were using combined oral contraceptive pills; one female had a non-hormonal (or copper) intrauterine device, and one had a contraceptive implant. A screening questionnaire was used to ensure the participants met the inclusion criteria and training/activity status. All participants were considered to be recreationally active as per the U.K. Chief Medical Officer’s guidelines of at least 150 min of moderate intensity exercise or 75 min of vigorous intensity exercise.

### Experimental Design

Participants visited the laboratory on two occasions. The first visit involved familiarisation with neuromuscular measures, followed by a treadmill incremental exercise test to the limit of tolerance. The second visit was the performance of a high-intensity interval exercise protocol, involving 4 × 3-min intervals at 90% of final incremental test velocity (v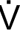
O_2peak_), interspersed with 90 sec rest periods. Before exercise, and in each rest period, participants performed a neuromuscular assessment while blood lactate was sampled, and heart rate and rating of perceived exertion (RPE) were recorded. Following exercise, the neuromuscular measures were repeated at 10, 20, and 30 min.

### Visit 1: Familiarisation & Incremental Exercise Testing

Participants were firstly familiarised with the neuromuscular stimulation techniques. This began with the determination of the femoral nerve stimulation threshold at rest (see details below), then the performance of warm up contractions increasing from 50% perceived effort to 90%. Participants then performed a neuromuscular function assessment (see details below).

Prior to the incremental exercise test, participants provided a capillary blood lactate sample from the earlobe, then performed a five-min warm up at an intensity corresponding to the first stage of the incremental test. Depending on each participant’s sex and training history, the speed for the first stage was set between 6 – 10 km·h^-1^. This permitted a similar number of subsequent stages to be completed prior to the limit of tolerance for males and females (6 ± 2 vs. 6 ± 1, respectively, *p* = 0.756). After the warmup, participants were given one minute rest, where another blood lactate sample was taken. The incremental test then involved three-min stages, with the treadmill speed increased by 1 km.h^-1^ each stage. Between stages, participants had 1 min of rest, where they provided further blood lactate samples. The treadmill was kept at a constant elevation of 1% throughout all testing and subsequent visits. Instructions were provided to “complete as many stages as possible”, and participants were informed that they should jump to the sides of the treadmill if they could not maintain the running speed any longer. The speed of the final complete stage was recorded as v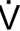
O_2peak_, however, if participants were able to run for longer than 90 secs during the final incomplete stage, 0.5 km.h^-1^ was added to the speed at the final complete stage when recording v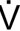
O_2peak_. Throughout the incremental exercise test, breath-by-breath gas exchange was recorded continuously, with heart rate and RPE (6-20 scale) recorded at the end of each stage.

Blood lactate samples were analysed immediately to determine lactate threshold and turnpoint (Biosen C-Line, EKF Diagnostics, Cardiff, UK). Lactate threshold was determined as the first work rate at which a non-linear increase in blood lactate concentration was observed, while lactate turnpoint was identified as the work rate that elicited a sudden and sustained increase in blood lactate concentration (Faude *et al*., 2009). The two lactate thresholds were analysed independently by two experimenters, and if any disagreement occurred, a third experimenter was consulted in order to mediate.

### Visit 2: High Intensity Interval Exercise

This visit began with the determination of the femoral nerve stimulation threshold, then warm up contractions increasing from 50% perceived effort to 90% were performed. Hereafter, a neuromuscular assessment was performed (see details below), before participants moved to the treadmill and provide a resting blood lactate sample.

Participants began the high-intensity interval exercise with a warmup consisting of a 5-min stage at 50% v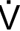
O_2peak_, and the final minute at 90% v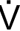
O_2peak_. After the warmup, participants rested for one min, while providing a blood lactate sample. The interval exercise involved 4 × 3-min intervals at 90% v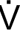
O_2peak_, interspersed with 90 secs of rest. At the end of each interval, participants moved from the treadmill to an isometric dynamometer, and performed three MVCs with femoral nerve stimulation, as well as providing a blood lactate sample, before returning to the treadmill. Heart rate and RPE were recorded at the end of each interval. Upon completion of the intervals, participants repeated the neuromuscular assessments immediately, as well as at 10-, 20-, and 30-min post-exercise. All participants completed the four intervals.

### Experimental Techniques Breath by Breath Gas Exchange

During all visits, expired gas was analysed breath-by-breath using an online system (Vyntus CPX, Jaeger, CareFusion, Germany). Oxygen (O_2_) and carbon dioxide (CO_2_) concentrations were analysed via a paramagnetic chemical fuel cell and non-dispersive infrared cell respectively. Before each test, the analysers were calibrated using ambient air and a gas of known O_2_ (15.00%) and CO_2_ (4.97%) concentrations. Ventilatory volumes were inferred from measurement of gas flow using a digital turbine transducer (volume 0 to 10 L, resolution 3 mL, flow 0 to 15 L·s^-1^), which was calibrated prior to each visit (Hans Rudolph Inc. Kansas City, USA).

### Neuromuscular Assessments

Pre-exercise neuromuscular assessments consisted of five MVCs separated by 30 secs. During the final three MVCs, femoral nerve stimulation was delivered at peak force of the MVC and 2 secs after. This was used to quantify MVC force, voluntary activation (VA), and potentiated twitch force (Q_tw.pot_). Following each interval, the same neuromuscular assessment was repeated but with three MVCs instead of five, as prior potentiation of twitches was not required (Kufel *et al*., 2002). For the assessments 10-, 20-, and 30-min post exercise, the baseline assessment was repeated including the two additional MVCs for potentiation of resting twitches.

### Femoral Nerve Stimulation

Electrical stimuli (200 *µ*s duration) were delivered to the femoral nerve via 32 mm-diameter surface electrodes (CF3200; Nidd Valley Medical, North Yorkshire, UK) using a constant-current stimulator (DS7AH, Digitimer, Welwyn Garden City, Hertfordshire, UK). The cathode was placed high in the femoral triangle over the nerve, and the anode was positioned midway between the greater trochanter and the iliac crest. The cathode was repositioned if necessary, to the location that elicited the largest quadriceps twitch amplitude (Q_tw_). Stimuli began at 20 mA, and increased by 20 mA until a plateau in Q_tw_ occured; this stimulus intensity was then increased by 30% to ensure supramaximal stimulations during the neuromuscular assessments.

### Force and Electromyography

For neuromuscular assessments, participants were seated on a custom-built MVC chair, with force (N) measured using a calibrated load cell (MuscleLab force sensor 300, Ergotest technology, Norway). The load cell was attached to the participant’s dominant leg, 2 cm superior to the ankle malleoli, using a non-compliant cuff. The load cell height was adjusted to ensure a direct line with the applied force for each participant. Participants were sat upright with knee and hip angles kept at 90° flexion. Force was sampled continuously (1000 Hz), and acquired for off-line analysis (Spike 2, Cambridge Electronic Design, Cambridge, UK).

### Blood Lactate Sampling

Blood lactate was sampled via capillary puncture technique with a 10 μl sample taken from the earlobe of each participant. Samples were immediately analysed for the concentration of lactate (mmol.L^-1^) and used for the calculation of lactate threshold and lactate turnpoint.

### Data Analysis

All MVCs were recorded, with the average of the peak forces during the three contractions used for further analyses. Voluntary activation assessed with nerve stimulation was calculated using the twitch interpolation method: VA (%) = (1 – [SIT/Q_tw,pot_] × 100), where SIT is the amplitude of the superimposed twitch force measured during MVC, and Q_tw,pot_ is the amplitude of the resting potentiated twitch force assessed 2 secs post-MVC.

Pulmonary gas exchange during both visits was exported in 5 secs bins, and the greatest 30 secs average recorded in visit 1 was used as peak values for each variable (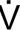
O_2_, 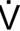
CO_2_, and 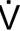
_E_). Data from visit two was exported in the same manner, with the final 30 secs of data during each interval expressed in absolute (L.min^-1^) and relative (%peak) units. Ventilatory equivalents of 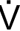
O_2_ and 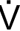
CO_2_ (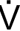
_E_/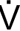
O_2_ and 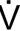
E/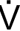
CO_2_) and respiratory exchange ratio (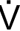
CO_2_/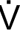
O_2_) were also calculated.

### Statistical Analysis

Data are presented as mean ± standard deviation within the text and figures. Normal distribution of data was confirmed with the Shapiro-Wilk test. As all variables had normally distributed data, males and females were compared with independent samples t tests for variables with a single value or time point. For repeated measures variables assessed during and after exercise, a two-way (sex × time) repeated measures ANOVA was performed. To assess fatigue during exercise, neuromuscular variables (normalised to % baseline) were assessed using a 2 × 5 (sex × time) ANOVA (time points: pre, interval 1, 2, 3, and 4). Whereas to assess recovery, neuromuscular variables (normalised to % baseline) were assessed with a 2 × 4 (sex × time) ANOVA (time points: post, +10, +20, and +30 min). Main and interaction effects were adjusted according to the Greenhouse-Geisser correction if the assumption of sphericity was violated, and significant effects were followed with Bonferroni-corrected *post-hoc* tests. The significance level for all statistical tests was set at α < 0.05.

## Results

### Incremental Exercise Testing

Absolute and relative data recorded during the incremental exercise test are presented in Table 1. As expected, males had greater values for absolute 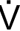
O_2peak_ (p < 0.001), however, when values were normalized to body mass, there was no sex difference in 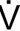
O_2peak_ (p = 0.292). Both sexes completed a similar number of incremental test stages before reaching exhaustion (p = 0.756). The velocity at which lactate threshold occurred, expressed as a percentage of v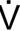
O_2peak_, was not different between sexes (males 67 ± 7% vs females 68 ± 9%, p = 0.773).

**Table 1:** Submaximal and peak data recorded during the incremental exercise test.

|  | Males (n = 10) | Females (n = 10) | P value |
| --- | --- | --- | --- |
| $\dot{V}O_{2\text{peak}}$ (l·min<sup>-1</sup>) | 3.901 ± 0.433 | 2.763 ± 0.323 | < 0.001 |
| $\dot{V}O_{2\text{peak}}$ (ml·kg<sup>-1</sup>·min<sup>-1</sup>) | 47.2 ± 6.0 | 44.4 ± 5.8 | 0.292 |
| $\dot{V}_{\text{Epeak}}$ (l·min<sup>-1</sup>) | 152.9 ± 20 | 105.8 ± 11.3 | < 0.001 |
| RER<sub>peak</sub> | 1.06 ± 0.05 | 1.05 ± 0.07 | 0.798 |
| HR<sub>peak</sub> (bpm) | 200 ± 4 | 200 ± 6 | 0.874 |
| $v\dot{V}O_{2\text{peak}}$ (km·h<sup>-1</sup>) | 14.9 ± 1.7 | 13.6 ± 1.6 | 0.090 |
| Lactate threshold (km·h<sup>-1</sup>) | 10.0 ± 1.8 | 9.2 ± 1.5 | 0.317 |

HR: heart rate; RER: respiratory exchange ratio; $\dot{V}_E$ : minute ventilation; $\dot{V}O_2$ : rate of oxygen uptake; $v\dot{V}O_{2\text{peak}}$ : velocity at the maximal rate of oxygen uptake.

### Metabolic & Cardiopulmonary Responses to Interval Exercise

During the interval task, heart rate, RPE, and blood lactate progressively increased (main effects of time: all p < 0.001). No main effects of sex were observed for heart rate (p = 0.601), RPE (p = 0.497) or blood lactate (p = 0.203). A sex × time interaction effect was observed for heart rate (Figure 1A, F_2.7, 45.5_ = 3.470, p = 0.028, ηp^2^ = 0.170), with *post-hoc* comparisons revealing females had greater heart rate at the end of the warmup (p = 0.041), but no other time points (p ≥ 0.696). No sex × time interaction effects were observed for either RPE (Figure 1B, p = 0.137) or blood lactate (Figure 1C, p = 0.183).

**Figure 1:**
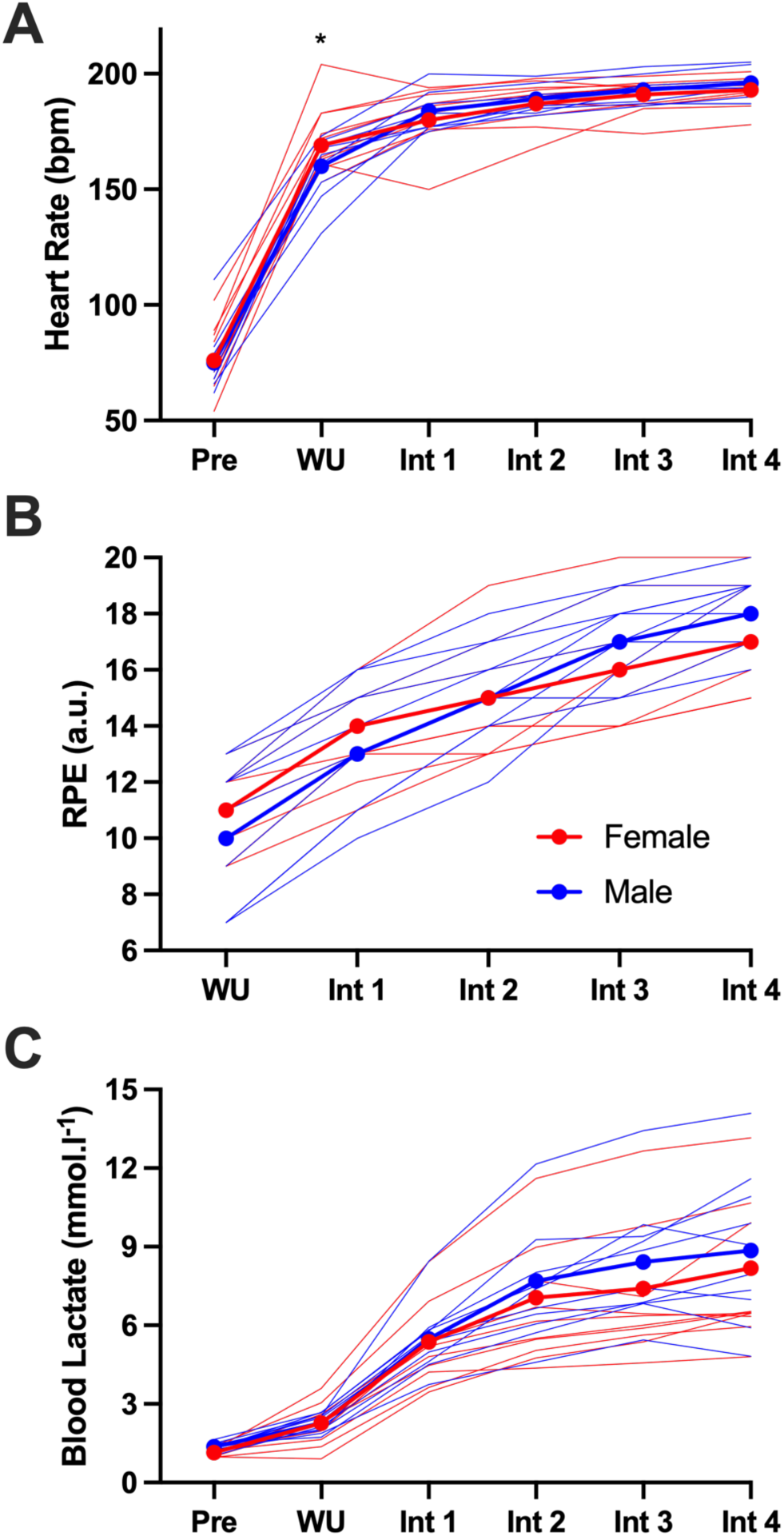
Heart rate (Panel A), rating of perceived exertion (RPE, Panel B), and blood lactate concentration (Panel C) at rest, following the warmup (WU), and at the end of each interval (Int). * = females greater than males (p < 0.05).

While males demonstrated greater absolute values (p < 0.001), when 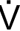
O_2_ and 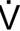
CO_2_ were expressed as a percentage of peak values (Figure 2A and 2B) from the incremental exercise test, no sex (p ≥ 0.631) or sex × time interaction effects (p ≥ 781) were observed. Similarly, no sex (p = 0.330) or sex × time interaction effects (p = 0.710) were observed for RER.

**Figure 2:**
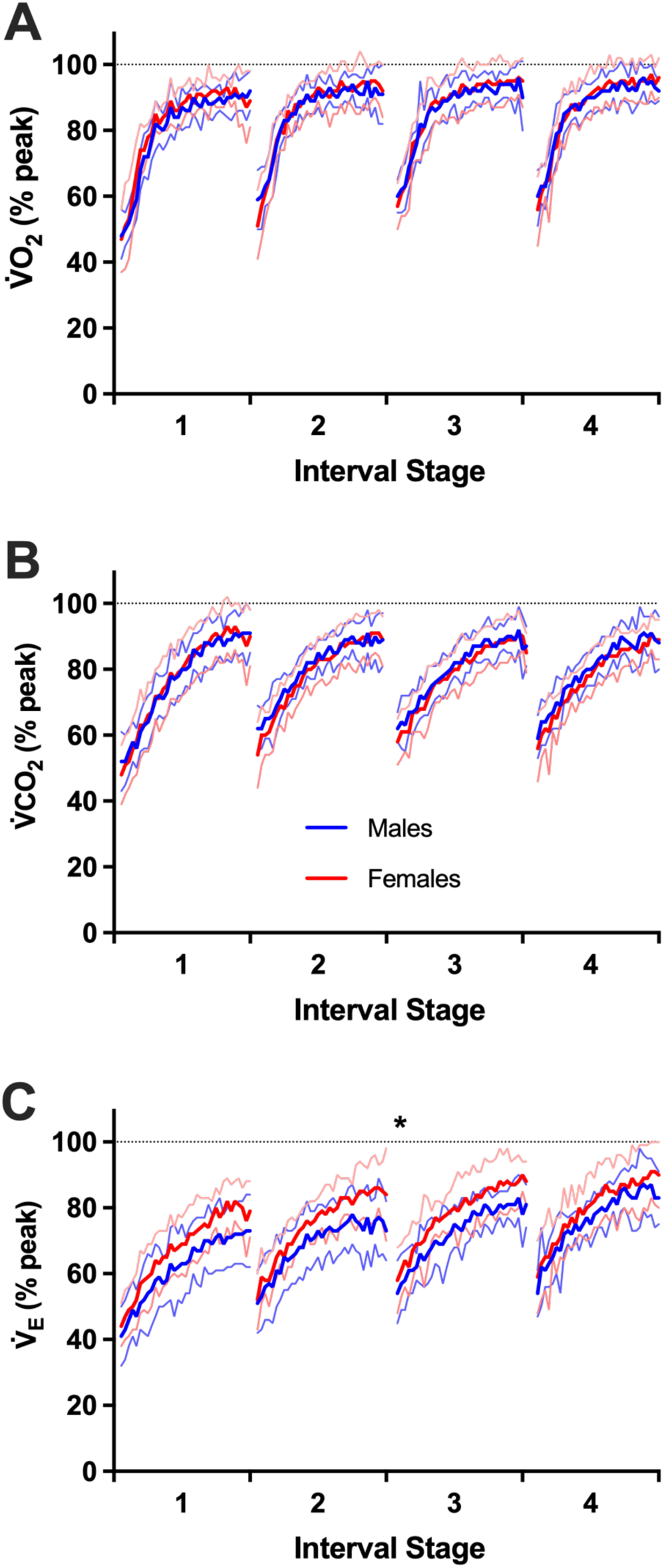
Rate of oxygen uptake (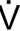
O2, Panel A), carbon dioxide production (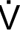
CO2, Panel B), and ventilation (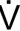
E, Panel C) during the four interval stages. Horizontal dashed lines indicate peak values attained during the incremental exercise test. * = sex x time interaction effect (p < 0.05).

In absolute values, males had greater 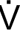
_E_ throughout the interval task (p < 0.001). When expressed as a percentage of peak values (Figure 2C), no main effect of sex was observed (p = 0.187), however, there was a sex × time interaction (F_3,54_ = 3.269, p = 0.042, ηp^2^ = 0.154) for 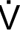
_E_. *Post-hoc* comparisons revealed no significant sex differences at any time point (p ≥ 0.050). Similarly, 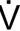
_E_/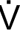
O_2_ demonstrated no main effect of sex (Figure 3A, p = 0.317), but a sex × time interaction effect (F_3,54_ = 4.831, p = 0.005, ηp^2^ = 0.212); with *post-hoc* comparisons revealing greater female values compared to males during the first interval (p = 0.034). 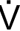
_E_/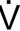
CO_2_ also demonstrated no main effect of sex (Figure 3B, p = 0.096), but a sex × time interaction effect (F_3,54_ = 2.853, p = 0.046, ηp^2^ = 0.137). *Post-hoc* comparisons revealed that females had greater values than males during the first interval for both variables (p ≤ 0.034) and second (p = 0.006) intervals for 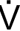
_E_/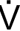
CO_2_.

**Figure 3:**
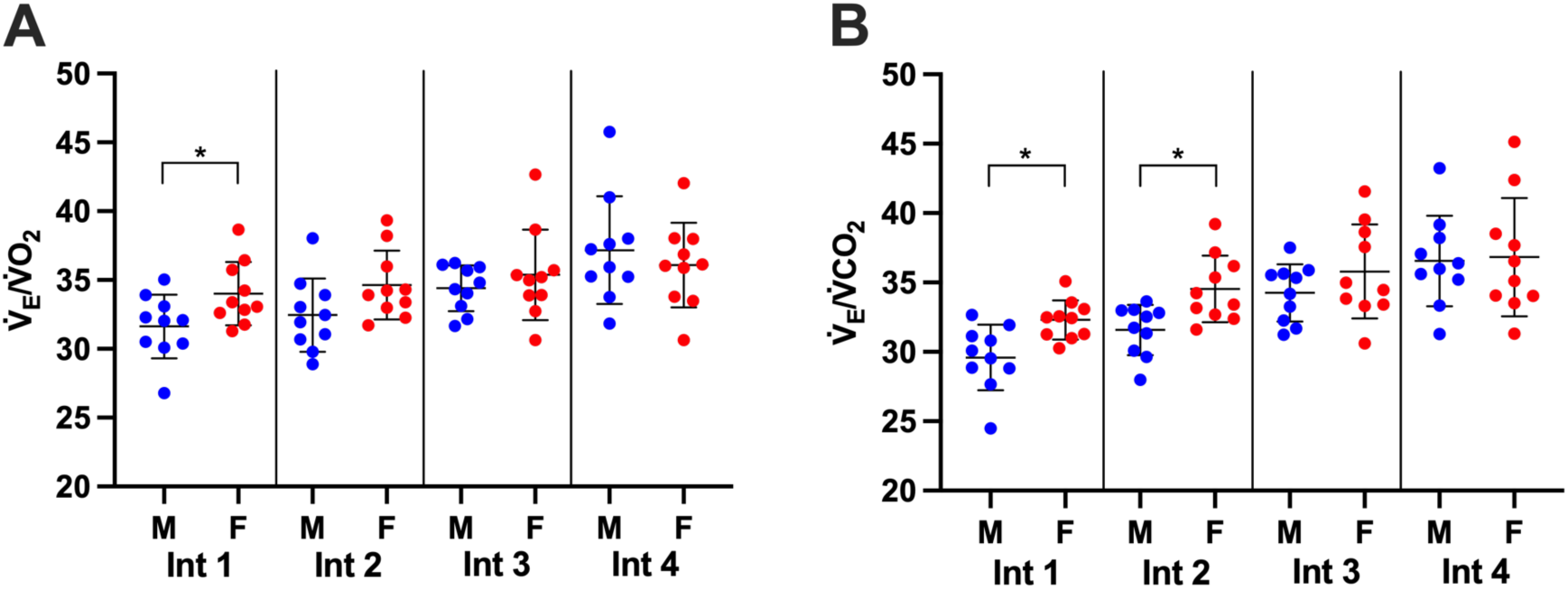
Ventilatory equivalents of oxygen uptake (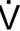
E/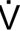
O2, Panel A) and carbon dioxide production (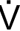
E/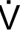
CO2, Panel B) during the final 30 secs of each interval (Int). Blue dots represent individual males, while red dots represent individual females. * = females greater than males (p < 0.05).

**Figure 4:**
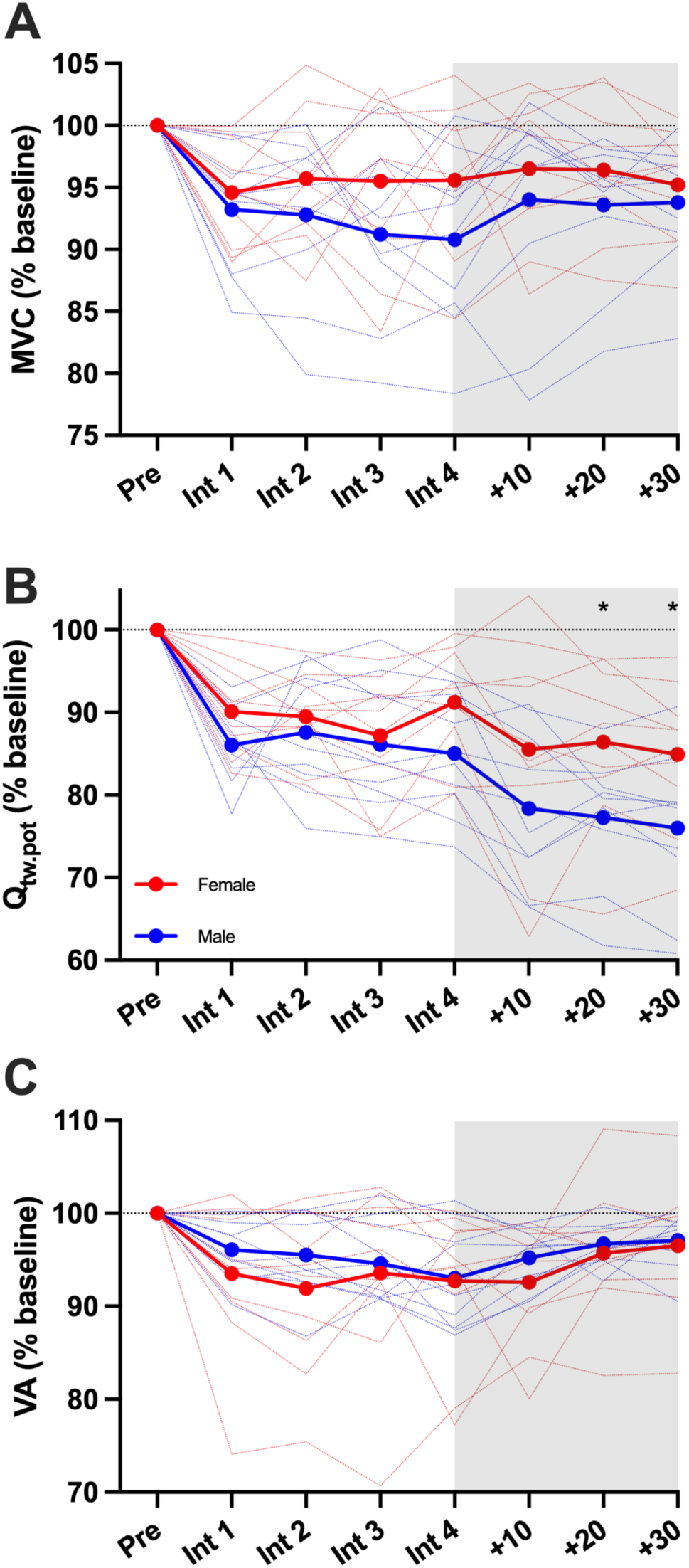
Neuromuscular variables recorded during and following the interval task. Thin dashed lines represent individual participants, whereas the solid lines represent group mean data. Maximal voluntary contraction (MVC, Panel A); quadriceps potentiated twitch (Qtw.pot, Panel B); and voluntary activation (VA, Panel C). * = females greater than males (p < 0.05).

### Fatigability and Recovery

At baseline, males produced greater maximal force than females (667 ± 35 vs. 471 ± 98 N, p < 0.001). Throughout the interval task, a main effect of time was observed for MVC (F_2.8,51.6_ = 9.897, p < 0.001, ηp^2^ = 0.355), Q_tw.pot_ (F_2.0,36.1_ = 30.481, p < 0.001, ηp^2^ = 0.629), and VA (F_2.4, 43.0_ = 10.884, p < 0.001, ηp^2^ = 0.377). *Post-hoc* comparisons revealed significant reductions in each variable after interval one compared to baseline (p ≤ 0.016), however, after the first interval, no further reductions were observed (p ≥ 0.506). No sex or sex × time interaction effects were observed for MVC (p ≥ 0.150), Q_tw.pot_ (p ≥ 0.184), or VA (p = 0.461).

In the 30-min recovery period after the interval task, no main effect of time was observed for MVC (p = 0.226), whereas as main effect of time was observed for Q_tw.pot_ (F_2.3, 42.0_ = 12.824, p < 0.001, ηp^2^ = 0.416) and VA (F_2.1,37.9_ = 6.531, p = 0.003, ηp^2^ = 0.266). No sex or sex × time interaction effects were observed for MVC (p ≥ 0.256) or VA (p ≥ 0.598), and while there was no sex × time interaction for Q_tw.pot_ (p = 0.567), there was a main effect of sex (F_1,18_ = 4.679, p 0.044, ηp^2^ = 0.206). *Post-hoc* comparisons indicated that females had greater Q_tw.pot_ amplitudes than males at 20 (p = 0.027) and 30 minutes (p = 0.030) after the interval task.

## Discussion

This study aimed to compare the physiological responses to high-intensity interval running exercise between sexes. It was hypothesised that there would be no sex differences in the metabolic response to the task, but females would be less fatigable. The lack of sex differences in the relative 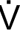
O_2_, RER, and blood lactate response to the task confirm the similarity of the metabolic response, whilst the sex difference in Q_tw.pot_ following the interval task indicated more fatigue-resistant female knee extensors. In addition, females demonstrated poorer ventilatory efficiency in the first half of the interval task compared to males, which was not evident in the second half of the task. Combined, these data demonstrate that the cardiopulmonary and neuromuscular responses to high-intensity interval running differ between sexes, adding to the growing evidence base that suggests practitioners should consider sex when prescribing exercise.

### Fatigability & Recovery Following Interval Exercise

The lack of sex difference in the decline in neuromuscular function during exercise contradicts previous literature utilising high-intensity constant load (Ansdell *et al*., 2020a; Azevedo *et al*., 2021; Azevedo *et al*., 2022) and self-paced (Solleiro Pons *et al*., 2023) locomotor exercise. Additionally, the data contradict those reported following repeated sprint exercise (Brooks *et al*., 1990; Billaut & Smith, 2009), reinforcing the notion that sex differences in fatigability are task-specific (Hunter, 2009). A recent study that employed a similar task to the present study (4 minute intervals with 3 minutes rest) also demonstrated similar fatigability between sexes (Tripp *et al*., 2024), suggesting that the specific demands of high-intensity interval exercise do not permit sex differences in fatigability from manifesting. The duration of the intervals used in the present study (3 min) likely resulted in a substantial metabolic disturbance within the working muscles. Jones *et al*. (2008) demonstrated that after 3.6 min of exercise at 110% of critical power, depletion of phosphocreatine (PCr) stores, inorganic phosphate accumulation, and pH decreases were all exaggerated compared to exercise below critical power. The rest periods between intervals (90 secs) likely permitted a degree of metabolic recovery of the working muscles, with near complete recovery of PCr stores and intramuscular pH being observed 90-120 secs after single-limb exercise (Iotti *et al*., 1991; Layec *et al*., 2013) and 6 minutes following all-out sprinting (Bogdanis *et al*., 1995). However, it should be acknowledged that the rest periods in the present study included the performance of MVCs, which may have delayed the metabolic recovery. The magnitude of knee-extensor fatigability that this repeated metabolic stress and recovery elicited was moderate, with MVC reductions (between 5-10%) less than was reported in males following a 5 km running time trial (Solleiro Pons *et al*., 2023), and substantially less than typically reported reductions following severe intensity cycling (∼20%, Thomas *et al.,* 2016; Brownstein *et al.,* 2021). One potential explanation why the fatigability induced by high-intensity interval running was not different between sexes is that the magnitude was too small to detect sex differences. Indeed, compared to previous data in single-limb and cycling exercise (Ansdell *et al*., 2020a; Azevedo *et al*., 2021; Azevedo *et al*., 2022), where sex differences were observed, the present study observed approximately half the degree of MVC reduction in both sexes.

Although sex differences in fatigability were not evident during the interval task, a sex difference was observed in the 30 min recovery period, whereby females demonstrated greater Q_tw.pot_ amplitudes relative to pre-exercise. Although this post-exercise period was termed the ‘recovery period’, no recovery was observed for either sex for MVC or Q_tw.pot_. Although central and peripheral contributions to fatigability are thought to mostly recover within 30 minutes following short duration exercise, recovery from running is further complicated by the presence of muscle damage caused by repeated stretch-shortening cycles (Carroll *et al*., 2017; Brownstein *et al*., 2021). Studies employing high- and low-frequency electrical stimulation to profile fatigue and recovery following damaging exercise reveal that low-frequency evoked contractions remain depressed, whereas high-frequency contractions recover (Skurvydas *et al*., 2016). Depression of low-frequency evoked contractions is thought to be underpinned by impaired intracellular calcium ion (Ca^2+^) release and/or reduced Ca^2+^ sensitivity of myofibrils (Bruton *et al*., 1998; Kamandulis *et al*., 2017), with the former being mechanistically linked to the reduced contractility of muscle fibres following damaging exercise (Gehlert *et al*., 2012). It is likely that the prolonged depression of MVC and Q_tw.pot_ in the present study was underpinned by altered Ca^2+^ handling within the knee-extensors, which also provides insight into the lesser relative reductions experienced by females compared to males. Evidence from Harmer *et al*. (2014) demonstrated sex differences in Ca^2+^ regulation before and after high-intensity exercise, with Ca^2+^ATPase activity reduced in males, but increased in females following exercise. Therefore, it is possible that the lesser relative reductions in Q_tw.pot_ experienced by females following high-intensity interval exercise in the present study were related to a lesser disruption to Ca^2+^ regulation within the knee-extensors. One additional consideration is that the patellar tendon stiffness has been demonstrated the be lower in females, which results in a greater mechanical buffer, and lesser knee-extensor fascicle lengthening for females during eccentric contractions (Hicks *et al*., 2013). Conceivably, this could also contribute to the lesser reductions in contractile function observed post-exercise in the present study.

### Metabolic and Cardiopulmonary Responses to Interval Exercise

The 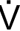
O_2_ recorded during all four intervals exceeded 90% of 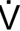
O_2peak,_ with values of ∼95% in both sexes by the final interval. The lack of sex difference in 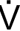
O_2_, 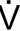
CO_2,_ RER, and blood lactate provides evidence that the metabolic response to high-intensity interval exercise was similar between males and females. This agrees with similar data recorded during constant-load exercise in the severe intensity domain, where no sex differences in the aforementioned variables were observed (Ansdell *et al*., 2020a). While sex differences in substrate utilisation have been observed previously, these appear to be limited to steady-state exercise (i.e., moderate and heavy intensity domains) rather than the intensities typically utilized during high-intensity interval exercise (Tarnopolsky *et al*., 1990; Devries *et al*., 2006). In response to the similar metabolic demands, males and females employed different respiratory strategies across the interval task, with females demonstrating poorer ventilatory efficiency during the first two intervals. Poorer ventilatory efficiency is thought to reflect poorer matching of lung ventilation to perfusion, and therefore impaired gas exchange (Sun *et al*., 2002). Sex differences are well-established in the structure and function of the respiratory system (Molgat-Seon *et al*., 2018), with larger airway diameters and lung volume, as well as greater alveolar surface area observed in males, even when height matched (Dominelli & Molgat-Seon, 2022). The lesser alveolar surface area in females results in poorer oxygen diffusing capacity during exercise (Bouwsema *et al*., 2017), necessitating greater relative 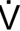
_E_ to achieve the same relative 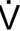
O_2_ and 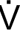
CO_2_. In addition to the present study that observed this sex difference during high-intensity interval exercise, greater relative 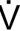
_E_ were observed in females during constant-load exercise in the heavy and severe intensity domains (Ansdell *et al*., 2020a). Interestingly, in the present study this ventilatory sex difference had dissipated by the third and fourth intervals, implying that both sexes ultimately experienced a loss of ventilatory efficiency as exercise progressed.

### Further Considerations

High-intensity interval exercise is often prescribed to improve an individual’s 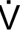
O_2max_ via positive adaptations to skeletal muscle capillary density, maximum stroke volume and cardiac output, and blood volume (MacInnis & Gibala, 2017). How sex influences physiological adaptation to exercise is unclear, with some evidence suggesting that 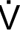
O_2max_ adaptation is blunted in females compared to males (Diaz-Canestro & Montero, 2019). Physiological adaptation to exercise is multi-factorial, with a variety of signalling pathways activated by disruptions to homeostasis in various physiological systems (Furrer *et al*., 2023). In the present study, the relative cardiovascular and metabolic perturbations were similar between sexes, but females experienced lesser contractile impairment following the task. Disruptions to intramuscular Ca^2+^ homeostasis that cause contractile impairment also activate Ca^2+^-calmodulin-dependent kinases (CaMK, Coffey & Hawley, 2007). CaMKs play an important role in regulating oxidative enzyme expression, as well as peroxisome proliferator-activated receptor gamma coactivator 1 (PGC-1a) and therefore mitochondrial biogenesis (Chin, 2005). While speculative, it could be suggested that the lesser reductions in contractile function experienced by females in the present study might result in a lesser stimulus for adaptation, however this hypothesis should be directly tested with appropriate methodologies.

### Limitations

The present study employed a high-intensity interval task that was performed at a percentage of each individual’s v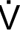
O_2peak_, which has previously been suggested to lead to greater heterogeneity of physiological responses compared to threshold-based approaches (Meyler *et al*., 2023). The approach of threshold-anchored exercise prescription has led to sex differences in fatigability and cardiopulmonary function being observed previously (Ansdell *et al*., 2020a; Azevedo *et al*., 2021; Azevedo *et al*., 2022). However, the present study observed similar sex differences when work rate was normalized to maximum capacity, rather than submaximal thresholds. Given that both sexes performed the task at ∼140% of their lactate threshold (Table 1), and reached 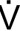
O_2_ values of ∼95% 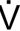
O_2peak_, peak blood lactate concentrations of ∼ 8 mmol·l^-1^, and peak RERs >1.05 during the interval task, it is likely that the task took place in the severe domain, and observed sex differences were not related to differences in the relative work rate of the task.

## Conclusions

This study demonstrated that males and females experienced similar cardiovascular and metabolic responses to a high-intensity interval exercise task, however females demonstrated poorer ventilatory efficiency in the first half of the task, and lesser reductions in knee-extensor contractile function following the task. Much like previous research that observed integrative sex differences during constant-load exercise, this study demonstrated that females and males do not experience the same responses to interval exercise. The exercise task employed in the present study is akin to those typically used for the enhancement of athletic performance, highlighting that those prescribing exercise should be cognisant that the sex of participants will influence the acute physiological responses.

## Acknowledgements

The researchers would like to thank the participants of the present study for their time and efforts.

## Reference

Ansdell P, Brownstein CG, Škarabot J, Hicks KM, Howatson G, Thomas K, Hunter SK & Goodall S. (2019a). Sex differences in fatigability and recovery relative to the intensity– duration relationship. The Journal of Physiology 597, 5577–5595.

Ansdell P, Brownstein CG, Škarabot J, Hicks KM, Simoes DCM, Thomas K, Howatson G, Hunter SK & Goodall S. (2019b). Menstrual cycle-associated modulations in neuromuscular function and fatigability of the knee extensors in eumenorrheic women. Journal of Applied Physiology 126, 1701–1712.

Ansdell P, Škarabot J, Atkinson E, Corden S, Tygart A, Hicks KM, Thomas K, Hunter SK, Howatson G & Goodall S. (2020a). Sex differences in fatigability following exercise normalised to the power–duration relationship. The Journal of Physiology 598, 5717–5737.

Ansdell P, Thomas K, Hicks KM, Hunter SK, Howatson G & Goodall S. (2020b). Physiological sex differences affect the integrative response to exercise: acute and chronic implications. Experimental Physiology 105, 2007–2021.

Ansdell P, Thomas K, Howatson G, Hunter S & Goodall S. (2017). Contraction intensity and sex differences in knee-extensor fatigability. Journal of Electromyography and Kinesiology 37, 68–74.

Azevedo RdA, Forot J, Iannetta D, MacInnis MJ, Millet GY & Murias JM. (2021). Slight power output manipulations around the maximal lactate steady state have a similar impact on fatigue in females and males. Journal of Applied Physiology 130, 1879–1892.

Azevedo RDEA, Forot J, Iannetta D, Aboodarda SJ, Millet GY & Murias JM. (2022). Time Course of Performance Fatigability during Exercise below, at, and above the Critical Intensity in Females and Males. Medicine and science in sports and exercise 54, 1665–1677.

Besson T, Parent A, Brownstein CG, Espeit L, Lapole T, Martin V, Royer N, Rimaud D, Sabater Pastor F, Singh B, Varesco G, Rossi J, Temesi J & Millet GY. (2021). Sex Differences in Neuromuscular Fatigue and Changes in Cost of Running after Mountain Trail Races of Various Distances. Med Sci Sports Exerc 53, 2374–2387.

Billaut F & Bishop DJ. (2012). Mechanical work accounts for sex differences in fatigue during repeated sprints. European Journal of Applied Physiology 112, 1429–1436.

Billaut F & Smith K. (2009). Sex alters impact of repeated bouts of sprint exercise on neuromuscular activity in trained athletes. Applied Physiology, Nutrition, and Metabolism 34, 689–699.

Boccia G, Dardanello D, Tarperi C, Festa L, La Torre A, Pellegrini B, Schena F & Rainoldi A. (2018). Women show similar central and peripheral fatigue to men after half-marathon*. European Journal of Sport Science 18, 695–704.

Bogdanis GC, Nevill ME, Boobis LH, Lakomy HK & Nevill AM. (1995). Recovery of power output and muscle metabolites following 30 s of maximal sprint cycling in man. The Journal of Physiology 482, 467–480.

Bouwsema MM, Tedjasaputra V & Stickland MK. (2017). Are there sex differences in the capillary blood volume and diffusing capacity response to exercise? Journal of Applied Physiology 122, 460–469.

Brooks S, Nevill ME, Meleagros L, Lakomy HKA, Hall GM, Bloom SR & Williams C. (1990). The hormonal responses to repetitive brief maximal exercise in humans. European Journal of Applied Physiology and Occupational Physiology 60, 144–148.

Brownstein CG, Millet GY & Thomas K. (2021). Neuromuscular responses to fatiguing locomotor exercise. Acta Physiologica 231, e13533.

Bruton JD, Lannergren J & Westerblad H. (1998). Mechanisms underlying the slow recovery of force after fatigue: importance of intracellular calcium. Acta Physiologica Scandinavica 162, 285–293.

Carroll TJ, Taylor JL & Gandevia SC. (2017). Recovery of central and peripheral neuromuscular fatigue after exercise. Journal of Applied Physiology 122, 1068–1076.

Chin ER. (2005). Role of Ca2+/calmodulin-dependent kinases in skeletal muscle plasticity. Journal of Applied Physiology 99, 414–423.

Coffey VG & Hawley JA. (2007). The Molecular Bases of Training Adaptation. Sports Medicine 37, 737–763.

Cowley ES, Olenick AA, McNulty KL & Ross EZ. (2021). “Invisible Sportswomen”: The Sex Data Gap in Sport and Exercise Science Research. Women in Sport and Physical Activity Journal 29, 146–151.

Crapo RO, Morris AH & Gardner RM. (1982). Reference values for pulmonary tissue volume, membrane diffusing capacity, and pulmonary capillary blood volume. Bull Eur Physiopathol Respir 18, 893–899.

Devries MC, Hamadeh MJ, Phillips SM & Tarnopolsky MA. (2006). Menstrual cycle phase and sex influence muscle glycogen utilization and glucose turnover during moderate-intensity endurance exercise. *American Journal of Physiology-Regulatory*, Integrative and Comparative Physiology 291, R1120–R1128.

Diaz-Canestro C & Montero D. (2019). Sex Dimorphism of VO2max Trainability: A Systematic Review and Meta-analysis. Sports Medicine 49, 1949–1956.

Diaz-Canestro C, Pentz B, Sehgal A & Montero D. (2022). Sex differences in cardiorespiratory fitness are explained by blood volume and oxygen carrying capacity. Cardiovascular Research 118, 334–343.

Dominelli PB & Molgat-Seon Y. (2022). Sex, gender and the pulmonary physiology of exercise. European Respiratory Review 31, 210074.

Dominelli PB, Render JN, Molgat-Seon Y, Foster GE, Romer LM & Sheel AW. (2015). Oxygen cost of exercise hyperpnoea is greater in women compared with men. The Journal of Physiology 593, 1965–1979.

Faude O, Kindermann W & Meyer T. (2009). Lactate Threshold Concepts. Sports Medicine 39, 469–490.

Furrer R, Hawley JA & Handschin C. (2023). The molecular athlete: exercise physiology from mechanisms to medals. Physiological Reviews 103, 1693–1787.

Gehlert S, Bungartz G, Willkomm L, Korkmaz Y, Pfannkuche K, Schiffer T, Bloch W & Suhr F. (2012). Intense Resistance Exercise Induces Early and Transient Increases in Ryanodine Receptor 1 Phosphorylation in Human Skeletal Muscle. PLOS ONE 7, e49326.

Georgescu VP, Thurston TS, Jarrett CL, Weavil JC, Richardson RS & Amann M. (2020). The female menstrual cycle: impact on cardiovascular, ventilatory and neuromuscular responses to whole body exercise. The FASEB Journal 34, 1–1.

Glace BW, Kremenic IJ & McHugh MP. (2013). Sex differences in central and peripheral mechanisms of fatigue in cyclists. European Journal of Applied Physiology 113, 1091–1098.

Glace BW, McHugh MP & Gleim GW. (1998). Effects of a 2-Hour Run on Metabolic Economy and Lower Extremity Strength in Men and Women. Journal of Orthopaedic & Sports Physical Therapy 27, 189–196.

Guenette JA, Querido JS, Eves ND, Chua R & Sheel AW. (2009). Sex differences in the resistive and elastic work of breathing during exercise in endurance-trained athletes. *American Journal of Physiology-Regulatory*, Integrative and Comparative Physiology 297, R166–R175.

Hammer SM, Sears KN, Montgomery TR, Olmos AA, Hill EC, Trevino MA & Dinyer-McNeely TK. (2023). Sex differences in muscle contraction-induced limb blood flow limitations. European Journal of Applied Physiology.

Harmer AR, Ruell PA, Hunter SK, McKenna MJ, Thom JM, Chisholm DJ & Flack JR. (2014). Effects of type 1 diabetes, sprint training and sex on skeletal muscle sarcoplasmic reticulum Ca2+ uptake and Ca2+-ATPase activity. The Journal of Physiology 592, 523–535.

Harms CA, McClaran SR, Nickele GA, Pegelow DF, Nelson WB & Dempsey JA. (1998). Exercise-induced arterial hypoxaemia in healthy young women. The Journal of Physiology 507, 619–628.

Hicks KM, Onambele-Pearson GL, Winwood K & Morse CI. (2013). Gender differences in fascicular lengthening during eccentric contractions: the role of the patella tendon stiffness. Acta Physiologica 209, 235–244.

Hunter SK. (2009). Sex Differences and Mechanisms of Task-Specific Muscle Fatigue. Exercise and Sport Sciences Reviews 37, 113–122.

Hunter SK. (2014). Sex differences in human fatigability: mechanisms and insight to physiological responses. Acta Physiologica 210, 768–789.

Hunter SK. (2016). Sex differences in fatigability of dynamic contractions. Experimental Physiology 101, 250–255.

Hunter SK, Critchlow A, Shin I-S & Enoka RM. (2004). Men are more fatigable than strength-matched women when performing intermittent submaximal contractions. Journal of Applied Physiology 96, 2125–2132.

Iotti S, Funicello R, Zaniol P & Barbiroli B. (1991). Kinetics of post-exercise phosphate transport in human skeletal muscle: An in vivo 31P-MR spectroscopy study. Biochemical and Biophysical Research Communications 176, 1204–1209.

James JJ, Klevenow EA, Atkinson MA, Vosters EE, Bueckers EP, Quinn ME, Kindy SL, Mason AP, Nelson SK, Rainwater KAH, Taylor PV, Zippel EP & Hunter SK. (2023). Underrepresentation of women in exercise science and physiology research is associated with authorship gender. Journal of Applied Physiology 135, 932–942.

Jones AM & Vanhatalo A. (2017). The ‘Critical Power’ Concept: Applications to Sports Performance with a Focus on Intermittent High-Intensity Exercise. Sports Medicine 47, 65–78.

Jones AM, Wilkerson DP, DiMenna F, Fulford J & Poole DC. (2008). Muscle metabolic responses to exercise above and below the “critical power” assessed using 31P-MRS. *American Journal of Physiology-Regulatory*, Integrative and Comparative Physiology 294, R585–R593.

Kamandulis S, de Souza Leite F, Hernández A, Katz A, Brazaitis M, Bruton JD, Venckunas T, Masiulis N, Mickeviciene D, Eimantas N, Subocius A, Rassier DE, Skurvydas A, Ivarsson N & Westerblad H. (2017). Prolonged force depression after mechanically demanding contractions is largely independent of Ca2+ and reactive oxygen species. The FASEB Journal 31, 4809–4820.

Kufel TJ, Pineda LA & Mador MJ. (2002). Comparison of potentiated and unpotentiated twitches as an index of muscle fatigue. Muscle & Nerve 25, 438–444.

Laursen PB & Jenkins DG. (2002). The Scientific Basis for High-Intensity Interval Training. Sports Medicine 32, 53–73.

Layec G, Malucelli E, Le Fur Y, Manners D, Yashiro K, Testa C, Cozzone PJ, Iotti S & Bendahan D. (2013). Effects of exercise-induced intracellular acidosis on the phosphocreatine recovery kinetics: a 31P MRS study in three muscle groups in humans. NMR in Biomedicine 26, 1403–1411.

MacInnis MJ & Gibala MJ. (2017). Physiological adaptations to interval training and the role of exercise intensity. The Journal of Physiology 595, 2915–2930.

Martin TR, Castile RG, Fredberg JJ, Wohl ME & Mead J. (1987). Airway size is related to sex but not lung size in normal adults. Journal of Applied Physiology 63, 2042–2047.

Mattu AT, Iannetta D, MacInnis MJ, Doyle-Baker PK & Murias JM. (2020). Menstrual and oral contraceptive cycle phases do not affect submaximal and maximal exercise responses. Scandinavian Journal of Medicine & Science in Sports 30, 472–484.

McDougall RM, Tripp TR, Frankish BP, Doyle-Baker PK, Lun V, Wiley JP, Aboodarda SJ & MacInnis MJ. (2023). The influence of skeletal muscle mitochondria and sex on critical torque and performance fatiguability in humans. The Journal of Physiology 601, 5295–5316.

Mead J. (1980). Dysanapsis in Normal Lungs Assessed by the Relationship between Maximal Flow, Static Recoil, and Vital Capacity. American Review of Respiratory Disease 121, 339–342.

Meyler S, Bottoms L, Wellsted D & Muniz-Pumares D. (2023). Variability in exercise tolerance and physiological responses to exercise prescribed relative to physiological thresholds and to maximum oxygen uptake. Experimental Physiology 108, 581–594.

Molgat-Seon Y, Peters CM & Sheel AW. (2018). Sex-differences in the human respiratory system and their impact on resting pulmonary function and the integrative response to exercise. Current Opinion in Physiology 6, 21–27.

Murphy WG. (2014). The sex difference in haemoglobin levels in adults — Mechanisms, causes, and consequences. Blood Reviews 28, 41–47.

Parker BA, Smithmyer SL, Pelberg JA, Mishkin AD, Herr MD & Proctor DN. (2007). Sex differences in leg vasodilation during graded knee extensor exercise in young adults. Journal of Applied Physiology 103, 1583–1591.

Parmar A, Jones T & Hayes P. (2021). The use of interval-training methods by coaches of well-trained middle- to long-distance runners. International Journal of Strength and Conditioning 1.

Roepstorff C, Donsmark M, Thiele M, Vistisen B, Stewart G, Vissing K, Schjerling P, Hardie DG, Galbo H & Kiens B. (2006). Sex differences in hormone-sensitive lipase expression, activity, and phosphorylation in skeletal muscle at rest and during exercise. American Journal of Physiology-Endocrinology and Metabolism 291, E1106–E1114.

Senefeld J, Pereira HM, Elliott N, Yoon T & Hunter SK. (2018). Sex Differences in Mechanisms of Recovery after Isometric and Dynamic Fatiguing Tasks. Med Sci Sports Exerc 50, 1070–1083.

Sheel AW, Dominelli PB & Molgat-Seon Y. (2016). Revisiting dysanapsis: sex-based differences in airways and the mechanics of breathing during exercise. Experimental Physiology 101, 213–218.

Sidhu SK, Cresswell AG & Carroll TJ. (2013). Corticospinal Responses to Sustained Locomotor Exercises: Moving Beyond Single-Joint Studies of Central Fatigue. Sports Medicine 43, 437–449.

Skurvydas A, Mamkus G, Kamandulis S, Dudoniene V, Valanciene D & Westerblad H. (2016). Mechanisms of force depression caused by different types of physical exercise studied by direct electrical stimulation of human quadriceps muscle. European Journal of Applied Physiology 116, 2215–2224.

Solleiro Pons M, Hunter SK & Ansdell P. (2023). Sex differences in fatigability and recovery following a 5 km running time trial in recreationally active adults. European Journal of Sport Science 23, 2349–2356.

Staron RS, Hagerman FC, Hikida RS, Murray TF, Hostler DP, Crill MT, Ragg KE & Toma K. (2000). Fiber Type Composition of the Vastus Lateralis Muscle of Young Men and Women. Journal of Histochemistry & Cytochemistry 48, 623–629.

Sun X-G, Hansen JE, Garatachea N, Storer TW & Wasserman K. (2002). Ventilatory Efficiency during Exercise in Healthy Subjects. American Journal of Respiratory and Critical Care Medicine 166, 1443–1448.

Tarnopolsky LJ, MacDougall JD, Atkinson SA, Tarnopolsky MA & Sutton JR. (1990). Gender differences in substrate for endurance exercise. Journal of Applied Physiology 68, 302–308.

Temesi J, Arnal PJ, Rupp T, Féasson L, Cartier R, Gergelé L, Verges S, Martin V & Millet GY. (2015). Are Females More Resistant to Extreme Neuromuscular Fatigue? Medicine and science in sports and exercise 47, 1372–1382.

Thomas K, Elmeua M, Howatson G & Goodall S. (2016). Intensity-Dependent Contribution of Neuromuscular Fatigue after Constant-Load Cycling. Med Sci Sports Exerc 48, 1751–1760.

Tripp Thomas R, Caswell Allison M, Aboodarda SJ & MacInnis Martin J. (2024). The Effect of Duration on Performance and Perceived Fatigability During Acute High-Intensity Interval Exercise in Young, Healthy Males and Females. Scandinavian Journal of Medicine & Science in Sports 34, e14692.

